# Diet of the Great Horned Owl *Bubo virginianus nigrescens* (Strigiformes: Strigidae) in the Northern Andes of Ecuador and a Literature Review of the Subspecies’ Dietary Patterns

**DOI:** 10.1101/2025.03.12.642946

**Authors:** Diego F. Cisneros-Heredia, Elias Viteri-Basso, Jorge Brito

## Abstract

Understanding the feeding ecology of top avian predators is crucial to unravelling trophic interactions and informing conservation strategies. Great Horned Owl *Bubo virginianus*, among the most widespread nocturnal raptors in America, remains understudied in the northern Andes. We investigated the diet of the subspecies *B. v. nigrescens* in an inter-Andean valley of northern Ecuador and compiled a literature review of its dietary patterns. Fifty-four pellets collected from 2014 to 2017 yielded 106 prey items. Mammals constituted the primary food source in frequency and biomass, with the Andean White-eared Opossum *Didelphis pernigra* and Black Rat *Rattus rattus* dominating the total biomass. Smaller native and introduced rodents were also common, while birds, bats, frogs, and Scarabaeid beetles appeared less frequently, underscoring the owl’s generalist foraging. A new record of predation on a young domestic cat highlights its adaptability to urbanised landscapes. Our literature review reveals similar trends across Colombia and Ecuador, with mammalian prey—often lagomorphs in higher elevations—dominant throughout. These findings suggest that *B. v. nigrescens* exhibits a flexible feeding ecology, capitalising on native and non-native prey. However, shifting land-use practices may affect prey availability and pose future challenges for this apex predator. Year-round and multi-elevation studies would help clarify seasonal variation and broader ecological dynamics, guiding conservation efforts in tropical mountain ecosystems.

## 1. Introduction

Top avian predators play key roles in structuring communities and regulating trophic dynamics. Analysing their dietary composition is essential to elucidate food-web relationships and responses to human-driven environmental change (Terraube and Bretagnolle, 2018). The Great Horned Owl *Bubo virginianus* is one of the most widespread and adaptable owl species in America, occupying a range of habitats from northern Canada to Uruguay and northeastern Argentina (Artuso et al., 2022; Fjeldså and Krabbe, 1990; Houston et al., 1998; Johnsgard, 1988; König and Weick, 2008; Mikkola, 2014; Olrog, 1968; Restall et al., 2006). Among its recognised subspecies, *Bubo virginianus nigrescens* inhabits the Andes of Colombia, Ecuador and northern Peru, yet its ecology remains less documented than that of its temperate North American counterparts (Artuso et al., 2022; Fjeldså and Krabbe, 1990; König and Weick, 2008; Restall et al., 2006; Traylor, 1958). Although many studies have characterised the broad diet of *B. virginianus* in North America—encompassing mammals, birds, reptiles, and invertebrates—data on the dietary habits of the South American populations are scarce (Aigner et al., 1994; Artuso et al., 2022; Bó et al., 2007; Bogiatto et al., 2003; Donázar et al., 1989; Freile et al., 2012; Houston et al., 1998; Houston and Duke, 2007; Johnsgard, 1988; Kremer and Belk, 2003; Restrepo-Cardona et al., 2019; Reynolds et al., 2021; Zimmerman et al., 1996). This gap hinders a fuller understanding of the species’ foraging ecology in tropical regions and highlights the need for targeted research.

Few studies have analysed the dietary composition of *B. virginianus* in heterogeneous Andean landscapes shaped by agriculture, urban development, and native vegetation across altitudinal gradients (Artuso et al., 2022; Cadena-Ortiz et al., 2022; Lehmann V., 1946, 1944; Pokines, 2007a). Documenting the diet of *B. v. nigrescens* in these areas would provide insights into the ecological processes that drive species interactions in mountain landscapes. Here, we record the dietary composition of a family of *B. v. nigrescens* in an inter-Andean basin of northern Ecuador, based on pellets collected over four years. Additionally, we offer a comprehensive literature review of the diet of the subspecies across its range. Our study contributes localised data that may inform ecological modelling, conservation planning, and broader assessments of the adaptive strategies that enable this owl to thrive in complex Andean settings.

## 2. Methodology

Fieldwork was carried out in the inter-Andean basin of Quito (also referred to as the *Hoya de Quito* or *Hoya de Guayllabamba*), a heterogeneous region in the northern Andes of Ecuador. This basin comprises the catchment area of the River Guayllabamba and includes multiple valleys and plateaus separated by gorges, ravines, and small hills (Alvarado et al., 2014; Cisneros-Heredia et al., 2015; Gondard, 1976; Stadel, 1991; Terán, 1984, 1982). This basin is largely isolated from other inter-Andean basins and the outer Andean slopes by mountains rising over 3000 m in elevation. Several major rivers flow through the basin, including the Pita, San Pedro, Machángara, Chiche, Pisque, and Monjas (Alvarado et al., 2014; Gondard, 1976; Stadel, 1991; Terán, 1984, 1982). Evidence of human occupation dates back to at least 10,000 years, and the city of San Francisco de Quito, Ecuador’s capital, was founded by Spanish colonisers in 1534 in its westernmost sector (Figueroa, 2017; Mothes et al., 1998; Torres Jiménez, 2024; Ugalde, 2019). Currently, much of the accessible terrain has been transformed into urban areas and agricultural lands, leaving only scattered forest remnants on steep slopes (Carrión and Erazo Espinosa, 2012; Terán, 1984).

*Bubo v. nigrescens* inhabits the highlands of Ecuador. It is considered most frequent in *páramo* landscapes above 3500 m and rarely found below this elevation (Chapman, 1926; Ridgely and Greenfield, 2001). Nonetheless, this owl has well-established yet dispersed populations in inter-Andean landscapes between 3100 and 2500 m, having adapted to living in anthropogenic habitats and likely recorded less frequently due to its inconspicuous behaviour (Cisneros-Heredia et al., 2015; Fjeldså and Krabbe, 1990; Lönnberg and Rendahl, 1922; McMullan and Navarrete, 2017; Salvadori and Festa, 1900; Stahl and Athens, 2001; Tellkamp, 2014). *B. v. nigrescens* is one of the largest raptors in the inter-Andean basin of Quito, where it is locally known as Cuscungo (Carrión, 1986; Ortiz Crespo, 1975; Wolf, 1892).

In 2014, a breeding pair of *B. v. nigrescens* (Fig. 1) was discovered inhabiting a mountain forest remnant in the canyon of the River Chiche (−0.193, −78.374, 2260 m elevation), approximately 11 km west of Quito, Pichincha province, Ecuador. From 2014 to 2017, pellets were collected once annually in December during Quito Christmas Bird Counts (Cisneros-Heredia et al., 2015). Each visit revealed one or two fledglings in the pair’s territory. All visible pellets—complete and fragmented—were gathered during a 30-minute search and stored individually in plastic bags.

**Figure 1.**
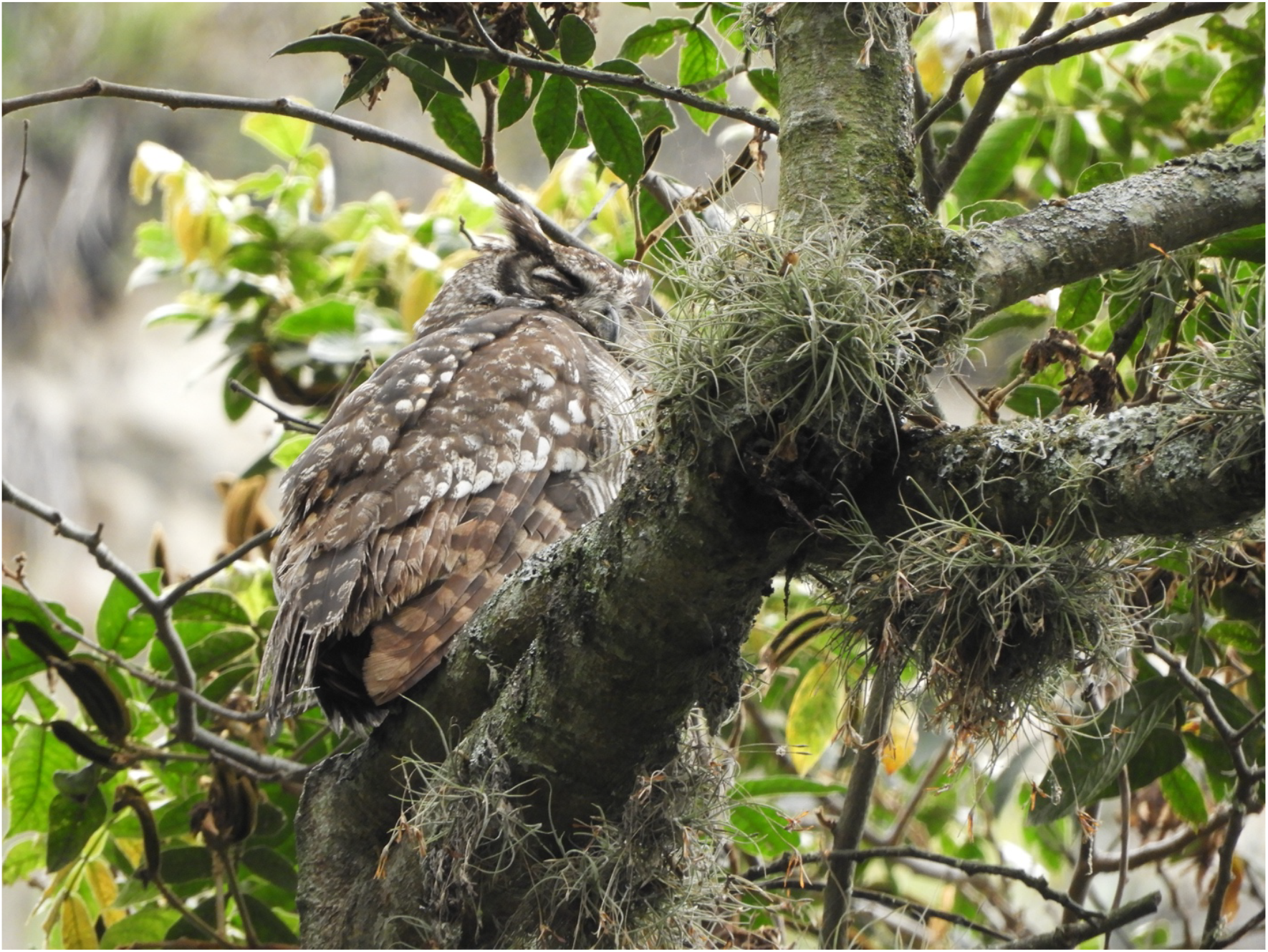
Great Horned Owl *Bubo virginianus* at the Canyon of the River Chiche. Photo by Elias Viteri-Basso.

In the laboratory, pellets were dried at room temperature and measured for length and width in millimetres (mm) using a digital calliper with a ±0.01 mm error margin (Buffalo® Tools). Weight was recorded on an analytical balance (capacity 120 g, precision 0.001 g, Fisher Scientific). Pellets were then dissected, their contents separated, labelled with unique locality codes, and deposited in the Instituto Nacional de Biodiversidad (MECN) collections, Quito, Ecuador. Prey items were identified using the reference collections of mammals, birds, reptiles, amphibians, and invertebrates at the MECN. The minimum number of individuals was determined by counting distinct skulls or calculating the greatest number of identical bones per taxon (Grayson, 2014; Lyman, 1994; Lyman et al., 2003; Mollhagen et al., 1972). Prey mass was estimated by multiplying the mean mass in grams (g) of each species by the corresponding number of individuals (Herrera and Jaksić, 1980). Across all samples, nearly all pellets contained at least one identifiable prey taxon. Disaggregated pellets often included fragments from multiple prey species that could not be assigned to a specific pellet number.

We conducted a comprehensive literature review of the diet of *B. v. nigrescens* using Google Scholar™ (https://scholar.google.com), employing the following search terms combined with Boolean operators: ‘Bubo’ AND ‘virginianus’ OR ‘nigrescens’ AND ‘diet’ AND ‘Ecuador’ OR ‘Colombia’ OR ‘Peru.’ We restricted localities to sites within the known range of *B. v. nigrescens* (Artuso et al., 2022; Houston et al., 1998; Traylor, 1958). We also searched in iNaturalist (http://www.inaturalist.org, California Academy of Sciences and National Geographic Society) and the Macaulay Library (https://www.macaulaylibrary.org, Cornell Lab of Ornithology) by specifying the species name and limiting results to the subspecies’ distribution; however, no relevant results were retrieved. We ran all searches on 9 February 2025 using on-site search engines.

## 3. Results

A total of 54 pellets were recovered between 2014 and 2017 (Table 1), yielding 106 individual prey items distributed among 16 species from three major vertebrate classes (Mammalia, Aves, and Amphibia) and one invertebrate order (Coleoptera). Mammals were the most significant dietary component, represented by three orders—Didelphimorphia, Rodentia, and Chiroptera (Fig. 2). Andean White-eared Opossum *Didelphis pernigra* and Black Rat *Rattus rattus* accounted for the highest biomass, whereas smaller rodents were captured more frequently but contributed less to the total biomass (Table 1). Juvenile *D. pernigra* consistently appeared as a dominant prey item across sampling years, suggesting elevated susceptibility of younger opossums to predation. Introduced and native rodents were common, highlighting the owl’s ability to exploit commensal and wild prey alike. Occasional captures of small bats further underscore the owl’s capacity to take a wide range of prey taxa.

**Table 1.** Diet of a family of Great Horned Owl *Bubo virginianus nigrescens*, based on pellets collected between 2014 and 2017, at a mountain forest remnant in the canyon of the River Chiche, Pichincha province, Ecuador.

| **CLASS** |  |  |  |  |  |
| --- | --- | --- | --- | --- | --- |
| **Order** | **Average** | **Number** | **%** | **Biomass** | **(%)** |
| **Family** | **mass** | **individuals** |  |  |  |
| **Species** |  |  |  |  |  |
| **MAMMALIA** |  |  |  |  |  |
| **Didelphimorphia** |  |  |  |  |  |
| **Didelphidae** |  |  |  |  |  |
| *Didelphis pernigra* | 400 | 22 | 20.8 | 8800 | 66.18 |
| **Rodentia** |  |  |  |  |  |
| **Cricetidae** |  |  |  |  |  |
| *Reithrodontomys soederstroemi* | 15 | 25 | 23.6 | 375 | 2.82 |
| *Akodon mollis* | 15 | 1 | 0.9 | 15 | 0.11 |
| *Phyllotis haggardi* | 20 | 4 | 3.8 | 80 | 0.60 |
| **Muridae** |  |  |  |  |  |
| *Mus musculus* | 14 | 2 | 1.9 | 28 | 0.21 |
| *Rattus rattus* | 162 | 18 | 17.0 | 2916 | 21.93 |
| *Rattus novergicus* | 210 | 1 | 0.9 | 210 | 1.58 |
| **Chiroptera** |  |  |  |  |  |
| **Phyllostomidae** |  |  |  |  |  |
| *Sturnira erythromos* | 16 | 2 | 1.9 | 32 | 0.24 |
| **Vespertilionidae** |  |  |  |  |  |
| *Eptesicus (fuscus) miradorensis* | 15.5 | 2 | 1.9 | 31 | 0.23 |
| **ANURA** |  |  |  |  |  |
| **Hemiphractidae** |  |  |  |  |  |
| *Gastrotheca riobambae* | 14 | 5 | 4.7 | 70 | 0.53 |
| **AVES** |  |  |  |  |  |
| **Columbiformes** |  |  |  |  |  |
| **Columbidae** |  |  |  |  |  |
| *Patagioenas fasciata* | 154 | 1 | 0.9 | 154 | 1.16 |
| **Passeriformes** |  |  |  |  |  |
| **Emberizidae** |  |  |  |  |  |
| *Zonotrichia capensis* | 22 | 4 | 3.8 | 88 | 0.66 |
| **Thraupidae** |  |  |  |  |  |
| *Rauenia bonariensis* | 25 | 1 | 0.9 | 25 | 0.19 |
| **Fringillidae** |  |  |  |  |  |
| *Spinus magellanicus* | 9 | 1 | 0.9 | 9 | 0.07 |
| **Turdidae** |  |  |  |  |  |
| *Turdus fuscater* | 130 | 3 | 2.8 | 390 | 2.93 |
| **Cardinalidae** |  |  |  |  |  |
| *Pheucticus chrysogaster* | 61 | 1 | 0.9 | 61 | 0.46 |
| **INSECTA** |  |  |  |  |  |
| **Coleoptera** |  |  |  |  |  |
| **Scarabeidae** | 1 | 13 | 12.3 | 13 | 0.10 |
| **Total** |  | **106** | **100.0** | **13297** | **100.00** |

**Figure 2.**
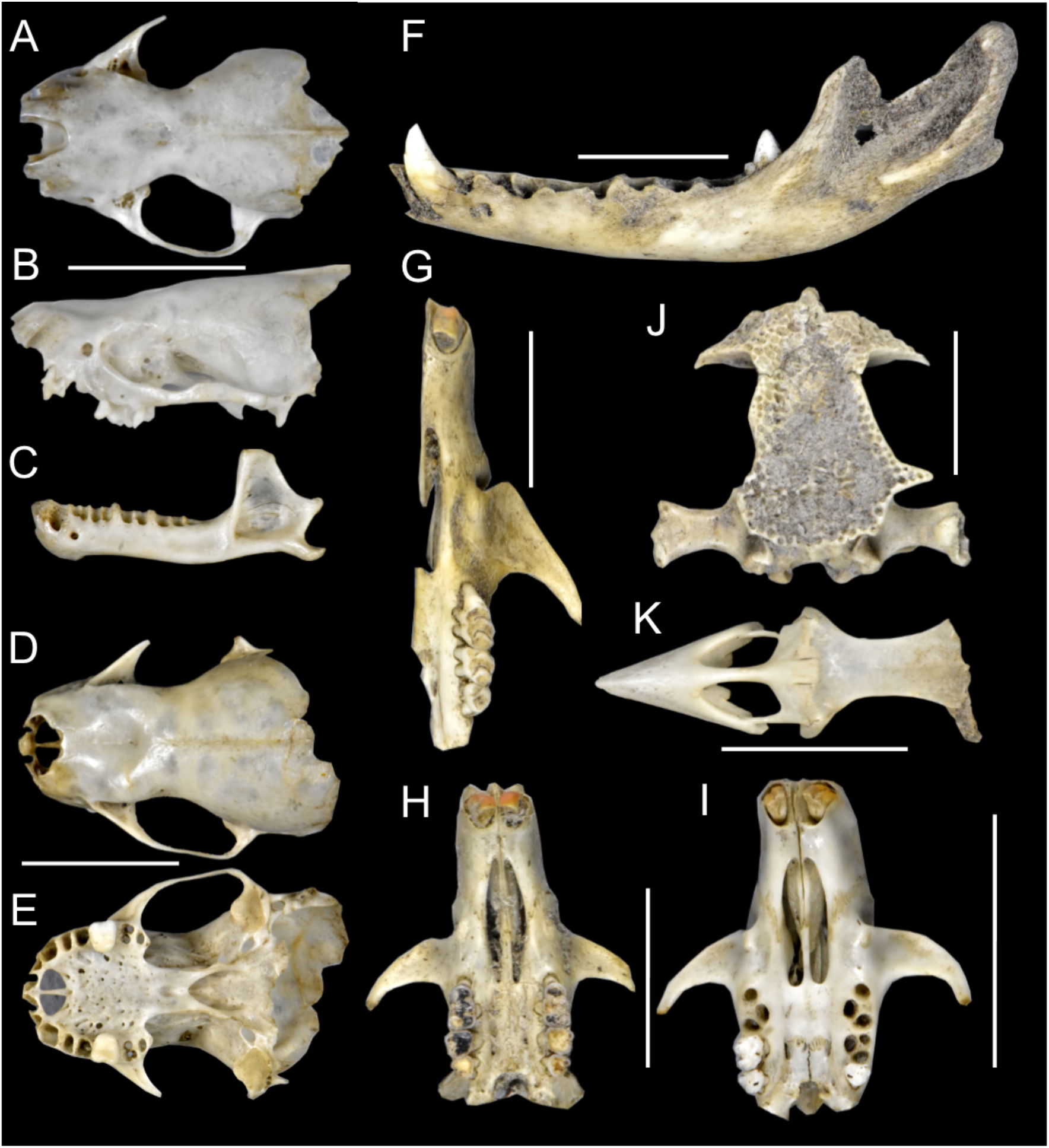
Some of the main owl prey in the River Chiche Canyon, Pichincha province, Ecuador. Dorsal (A), lateral (B) and mandible (C) views of the skull of *Eptesicus miradorensis*; dorsal (D) and ventral (E) views of the skull of *Sturnira erythromos*; lateral view of the mandible (F) of a juvenile of *Didelphis pernigra*; partial ventral view (G) of the skull of *Rattus rattus*; ventral view (H) of the skull of *Phyllotis haggardi*; ventral view (I) of the skull of *Reithrodontomys soederstroemi*; part of the beak and skull (K) of *Zonotrichia capensis*; and dorsal view of the skull (J) of *Gastrotheca riobambae*. Scale bars = 10 mm. Photos by Jorge Brito.

Birds collectively constituted a smaller proportion of the total biomass, yet they included a diverse array of species, ranging from medium-sized doves to smaller passerines. Amphibians were detected at a low frequency, represented solely by Quito Marsupial Frog *Gastrotheca riobambae* (Table 1) Scarabaeid beetles (Coleoptera) were the sole invertebrates identified and contributed minimally to the total biomass. These findings demonstrate the Great Horned Owl’s preference for a mammal-based diet while showing its ability to use a generalist and opportunistic feeding strategy in the inter-Andean basin of Quito.

Our literature review yielded documented prey records for *B. v. nigrescens* at multiple sites in Colombia and one in Ecuador (Table 2). No published data were located for northern Peru. Most studies used direct field observations, pellet analyses, or nest remains to identify prey. They generally consist of short natural history accounts with specific geographic or temporal scopes, with few providing extensive dietary analyses. Early references often presented anecdotal or observational accounts, while recent works employed systematic approaches and offered more quantifiable evidence. These studies illustrate that *B. v. nigrescens* is a resourceful predator with a broad taxonomic reach and opportunistic predation habits. Mammalian prey dominates in frequency and biomass across all studied sites, especially lagomorphs, rodents and opossums. In one study at Tolima (Colombia), Andean Cottontail *Sylvilagus andinus* accounted for most prey items (Restrepo-Cardona et al., 2019). Small native and non-native rodents appear regularly in the diet of *B. v. nigrescens*, highlighting the importance of small mammals in the owl’s diet.

**Table 2.**
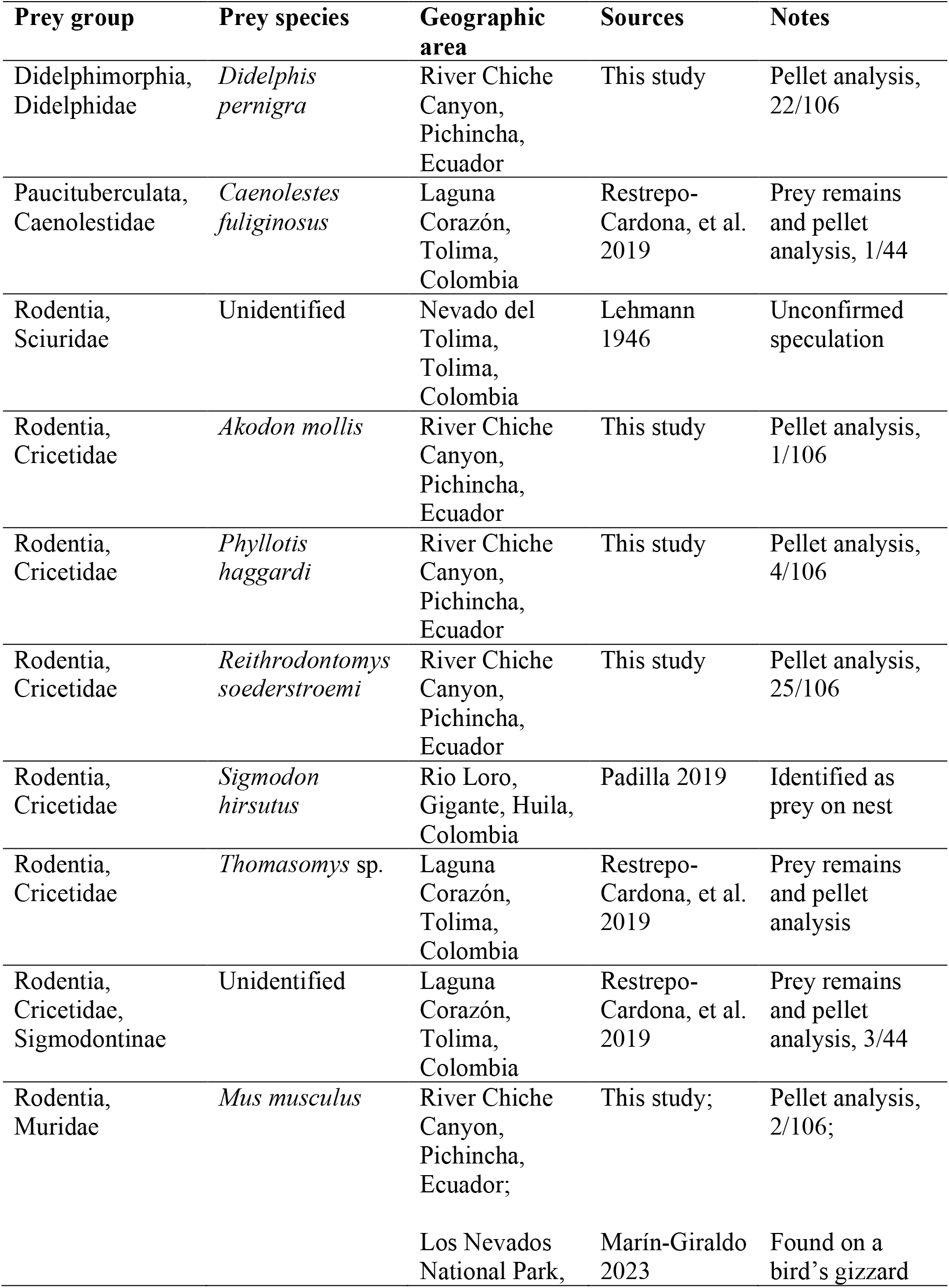
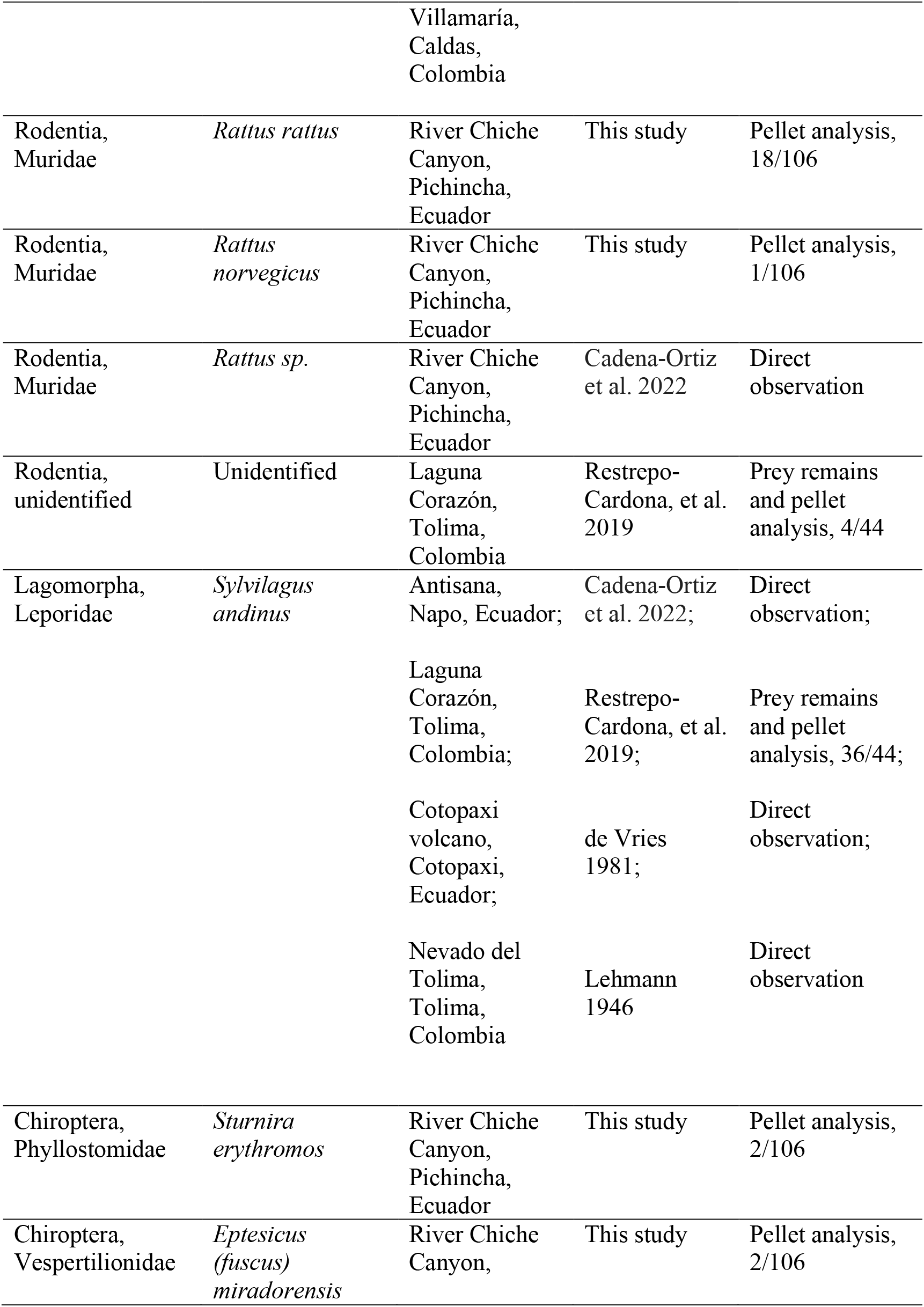
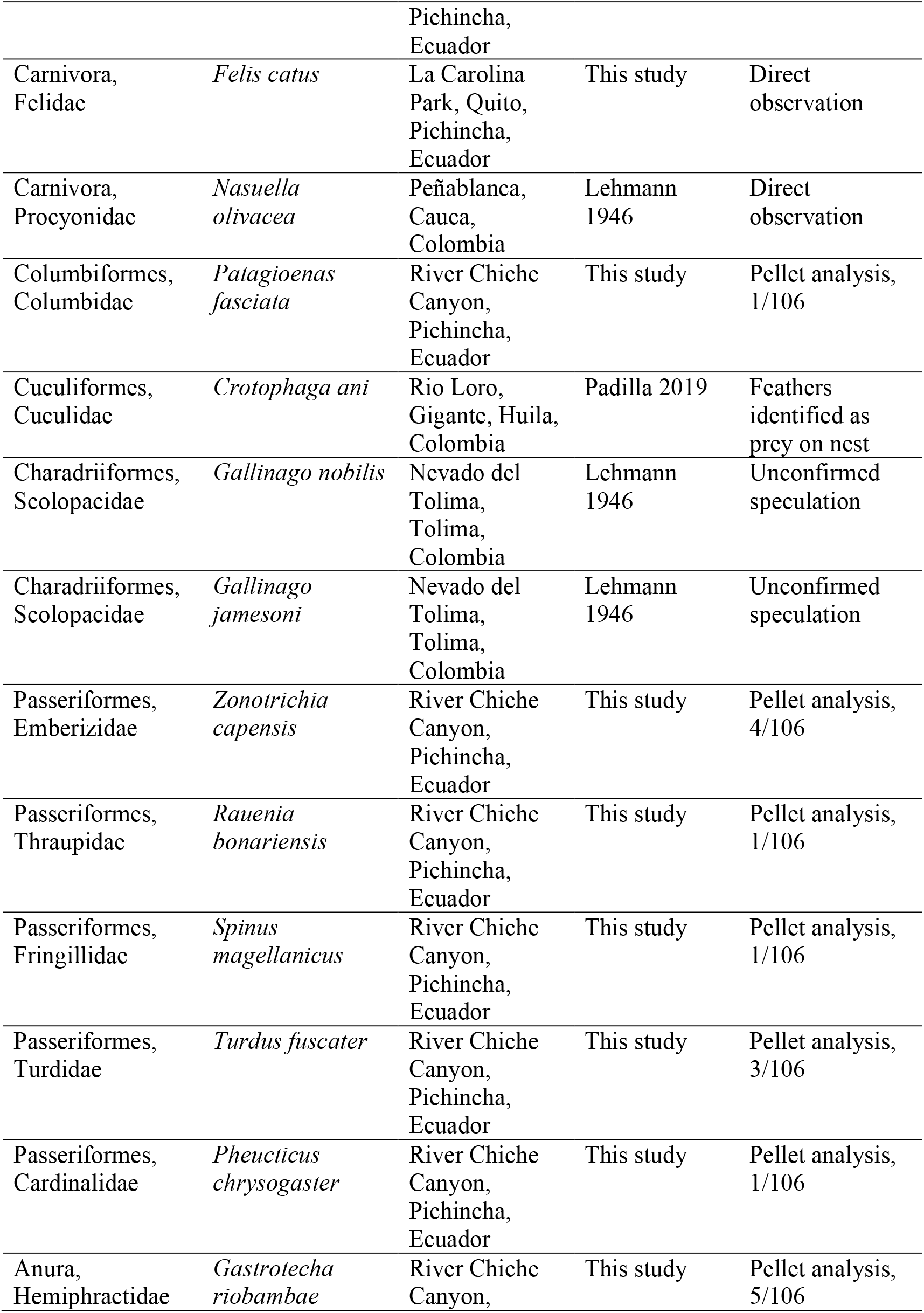
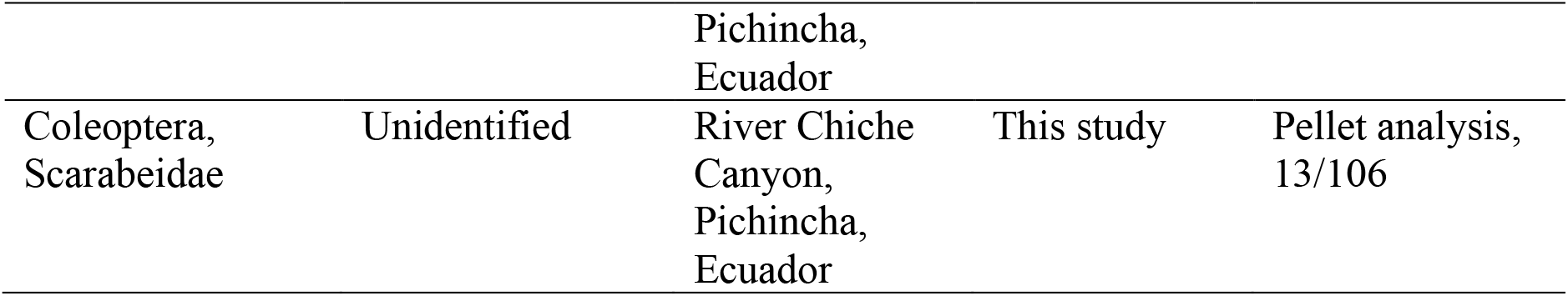
List of known prey of Great Horned Owl *Bubo virginianus nigrescens*.

Occasionally, *B. v. nigrescens* captures larger prey, such as Mountain Coati *Nasuella olivacea* (Lehmann V., 1946, 1944). Additionally, one of the authors of this study observed an adult *B. v. nigrescens* eating a young, feral House Cat *Felis catus* in a large urban park (La Carolina Park) in Quito, Ecuador (D. F. Cisneros-Heredia, pers. obs. February 2001).

## 4. Discussion

Our analysis of *Bubo virginianus nigrescens* pellets from the inter-Andean basin of Quito reveals a broad and flexible feeding strategy within a landscape increasingly influenced by agriculture and urban development. Mammals dominated the diet in frequency and biomass, with medium-sized opossums (*Didelphis pernigra*) and small rodents as the most frequent prey. This pattern parallels trends observed in other temperate and tropical populations of Great Horned Owl (Aigner et al., 1994; Artuso et al., 2022; Bogiatto et al., 2003; Cadena-Ortiz et al., 2022; Dias and Kasper, 2023; Donázar et al., 1989; Houston et al., 1998; Houston and Duke, 2007; König and Weick, 2008; Kremer and Belk, 2003; Mikkola, 2014; Osses and Bernardis, 2018; Pokines, 2007b; Restrepo-Cardona et al., 2019; Reynolds et al., 2021; Tomazzoni et al., 2004; Zimmerman et al., 1996). Notably, *Didelphis pernigra* featured prominently in our samples, with juveniles making up a substantial proportion of the opossum remains. Younger individuals may be more susceptible to predation due to greater availability, limited defensive behaviours, or a combination of factors. Comparable findings were reported by Wink et al. (1987), who recorded Virginia Opossum *Didelphis virginiana* as the primary biomass contributor in Great Horned Owl diets in Pennsylvania.

Although large prey items are less commonly documented, Great Horned Owls are known to capture animals up to the size of young foxes, house cats, and skunks (Houston et al., 1998). Our observation of a predation event on a young house cat underscores the owl’s occasional targeting of non-wild prey, revealing potential human-wildlife interactions as domestic animals increasingly overlap with avian predators (Washburn, 2016). The relatively high prevalence of introduced rodents in our samples further illustrates the owl’s capacity to exploit anthropogenic landscapes, taking advantage of commensal non-native rodents that often thrive near human settlements and agricultural areas (Cadena-Ortiz et al., 2022; Houston et al., 1998; Osses and Bernardis, 2018; Wink et al., 1987). In contrast, avian prey tends to be recorded less frequently. However, avian prey may be underrepresented in pellet analyses, which often produce fewer identifiable bird remains (Woodman et al., 2005). Nonetheless, predation on birds, bats and insects confirms the owl’s extensive dietary breadth.

Our record of two bat specimens in pellets from December 2015 adds a fourth locality in Ecuador for the Brown Bat *Eptesicus* (*fuscus*) *miradorensis* (Chiroptera: Vespertilionidae) (Fig. 2). This species was previously documented only at three inter-Andean sites (“above Quito, 11 feet altitude [=3350 m]”, Lönnberg 1921; River Pisque and Quebrada Trapichuco; Arguero and Albuja 2012) (Ramírez-Chaves et al. 2023). Such bat occurrences highlight the broader value of raptor diet studies for advancing knowledge of poorly known mammal distributions (Cadena-Ortiz et al., 2023; Heisler et al., 2016; Stutz et al., 2020).

In most Andean regions, cottontail rabbits contribute substantially to the Great Horned Owl’s diet biomass (De Vries, 1981; Lehmann V., 1946; Restrepo-Cardona et al., 2019). However, they were absent from our site. This absence may reflect habitat preferences, seasonal availability, or competition with other predators. *Sylvilagus andinus* typically inhabits mountainous areas between 3000 and 4000 m elevation but descends into lower open valleys and plateaus within inter-Andean basins (Beltrán-Ortiz et al., 2017; Cadena-Ortiz et al., 2019; Hershkovitz, 1938; Reina Moreno, 2019; Trujillo G. and Trujillo G., 2007). However, it appears to avoid canyons and gorges, where the owl family from our sampling most probably forages, thus reducing the likelihood of encounters. Nearby, at the Tababela plateau a few kilometers from our study site, *S. andinus* has been documented in the diets of American Barn Owl *Tyto furcata* and Andean Fox *Lycalopex culpaeus* (Cadena-Ortiz et al., 2019; Reina Moreno, 2019). It is also worth noting that we collected pellets only in December each year, so we cannot rule out seasonal or interannual shifts in prey composition, including changes in rabbit activity outside our sampling window.

Consistent with our findings, the literature review depicts *B. v. nigrescens* as a resourceful generalist predator across the northern Andes, capitalising on the most readily available mammals, usually focusing on one or two profitable species within the region (Hindmarch and Elliott, 2014; Houston et al., 1998; Llinas-Gutiérrez et al., 1991; Marchesi et al., 2002), while also including a wide range of vertebrates and occasional insects (Cadena-Ortiz et al., 2022; De Vries, 1981; Lehmann V., 1946, 1944; Marín-Giraldo et al., 2023; Padilla, 2019; Restrepo-Cardona et al., 2019). Such dietary flexibility likely underpins its resilience in a landscape with significant anthropogenic modifications (Artuso et al., 2022; Houston et al., 1998; Houston and Duke, 2007). Mammalian prey generally dominates the diet of *B. v. nigrescens* across the northern Andes (Cadena-Ortiz et al., 2022; De Vries, 1981; Lehmann V., 1946, 1944; Marín-Giraldo et al., 2023; Padilla, 2019; Restrepo-Cardona et al., 2019). Small rodents and lagomorphs account for nearly 90% of the Great Horned Owl’s diet biomass in temperature regions of North and South America (Donázar et al., 1989; Houston et al., 1998; Knight and Jackman, 1984; Kremer and Belk, 2003; Llinas-Gutiérrez et al., 1991; Teta, 2006; Zimmerman et al., 1996). In contrast, in the Neotropics, the species has a more diverse diet, with mammals still playing a significant role but with a higher proportion of birds, comprising 15% to 35% of the total biomass, and including a greater proportion of amphibians and reptiles (Dias and Kasper, 2023; Pokines, 2007b; Tomazzoni et al., 2004).

The Great Horned Owl’s adaptability may help it remain a top predator in novel ecosystems. However, ongoing, accelerated land-use changes across the Andes may alter prey communities by favouring invasive rodents and constraining native species. Such shifts raise concerns about ecological balance, potential disease transmission, and the long-term viability of native fauna. Future investigations, including year-round sampling and comparative studies across diverse elevations or degrees of urbanisation, would provide deeper insights into *B. v. nigrescens* prey selection, population trends, and its overarching role in Andean food webs.

## Acknowledgements

We are grateful to Juan Manuel Carrión for introducing us to the area inhabited by the owl family, to María Elena Heredia and Jonathan Guillemot for their valuable companionship and support during fieldwork, and to Glenda Pozo (Instituto Nacional de Biodiversidad INABIO) for identifying bird remains in the pellets. This research was possible thanks to Colectivo AvesQuito, which coordinates the Quito Christmas Bird Counts; INABIO, which provided access to its scientific collections for prey identification; the Biodiversity Heritage Library, which offers open-access literature; and the Universidad San Francisco de Quito USFQ, Instituto de Biodiversidad Tropical for their support during lab work.

## CRediT authorship contribution statement

**Diego F. Cisneros-Heredia:** Conceptualisation, Methodology, Validation, Formal analysis, Investigation, Resources, Data curation, Writing - original draft, Writing - review & editing, Visualization, Supervision, Project administration, Funding acquisition. **Elias Viteri-Basso:** Validation, Investigation, Data curation, Writing - original draft, Writing - review & editing. **Jorge Brito:** Methodology, Validation, Formal analysis, Investigation, Resources, Data curation, Writing - original draft, Writing - review & editing, Visualization, Funding acquisition.

